# Exploring local and regional drivers of microbial biodiversity across freshwater ponds

**DOI:** 10.64898/2025.12.16.694477

**Authors:** Emily A. Hardison, José Goyco-Blas, Noah Leith, Trina M. Wantman, Jakub Zegar, Matthew W. H. Chatfield, Michel Ohmer, Kevin D. Kohl

## Abstract

**Aim:** Ponds are isolated, highly variable environments that exhibit low spatial autocorrelation of environmental variables and have island-like features, which may give rise to unique biogeographic patterns compared to terrestrial and lotic environments. Here, we evaluate whether commonly observed biogeographical patterns apply to microbial biodiversity in pond fungal and bacterial communities. Specifically, we tested (1) whether these communities follow latitudinal diversity gradients and distance decay relationships, and (2) if variation in community composition or richness was related to specific environmental or land-use factors.

**Location:** Eastern, USA

**Time period:** Summer, 2022

**Taxa:** Bacteria and Fungi

**Methods:** We collected water and muck from 39 ponds across 8 states in the Eastern USA. We extracted DNA from these samples and sequenced sections of the 16S rRNA and ITS1 genes to survey the bacterial and fungal communities, before evaluating biogeographical patterns.

**Results:** We found evidence of latitudinal diversity gradients in muck fungal communities, but not bacterial communities. We observed weak distance decay relationships in all sample types. Community richness was related to some environmental filters, where conductivity was positively related to water bacterial richness, but negatively related to muck fungal richness. Environmental drivers explained low to moderate variation in microbial composition, with temperature universally linked to microbial biodiversity.

**Conclusions:** Pond microbiomes exhibit unique biogeographic patterns depending on the microhabitat and microbial taxa in question, with land use and abiotic conditions, especially temperature, explaining some variation in microbial biodiversity across sites. Our findings suggest that the low spatial autocorrelation in environmental conditions and the lack of connectivity across ponds provides a useful framework for investigating localized drivers of microbial diversity.

## Introduction

Freshwater ecosystems occupy a small fraction of Earth’s surface, yet harbor a disproportionally high percent of global biodiversity (Dudgeon *et al*., 2006; Strayer & Dudgeon, 2010). Among freshwater systems, ponds are often overlooked, despite being biodiversity hotspots which have small catchments and interface closely with human activity (Hill *et al*., 2021). They are also obligate habitat for several species that exhibit high site fidelity. Ponds exist across a gradient of urban and natural landscapes, including pristine naturally occurring wetlands, human-made decorative yard features, and aquaculture grow-out facilities (Veach *et al*., 2021). Within these environments, microbes living in the pond water and substrates serve as important regulators of biogeochemical cycles (Grossart *et al*., 2020; Premke *et al*., 2022) and a source pool from which many pond-dwelling plants and animals acquire their host-associated microbiomes (Rudzki et al. 2026; Jones et al., 2025). Thus, these communities are essential to maintaining a healthy and diverse pond ecosystem. However, the factors shaping microbial diversity across regional or global scales have been understudied compared to biogeographic patterns in the diversity of plants and animals (Astorga *et al*., 2012), especially in unique and understudied environments, like ponds.

Latitudinal diversity gradients, whereby taxonomic richness tends to increase with decreasing latitude, are one of the most pervasive large-scale biodiversity trends (Hillebrand, 2004). This trend has been observed across several taxonomic groups and environments, though strength of the gradient differs across environments, studies, and taxa (Kinlock *et al*., 2018). For example, a recent meta-analysis by Kinlock et al., [2018], found that bacteria and fungi did not show consistent latitudinal diversity gradients, nor did freshwater environments overall (across taxa). In contrast, others have found evidence of latitudinal diversity gradients in inland water bacterial communities (Wang *et al*., 2024) and some fungal soil communities (Tedersoo *et al*., 2014). Differences across studies may reflect unique mechanisms acting on microbial biodiversity across environments and taxa. Several mechanisms have been proposed to govern latitudinal diversity gradients in macro-organisms, including environmental gradients, strength of species interactions, dispersal limitations that co-vary with latitude (Hillebrand, 2004; Huang *et al*., 2024). However, more research is needed to resolve whether latitudinal diversity patterns apply to microbes across various environments along with the primary processes driving broad spatial gradients in microbial diversity.

Distance is generally a key driver of spatial variation in community composition, though the effects of distance are thought to differ between micro and macro-organisms. In macro-organism ecology, the farther apart sites are, the more environmental conditions and dispersal are expected to limit overlaps in membership. This leads to a ‘distance decay’ relationship, where the similarity in species composition between sites decreases the farther apart sites are (Astorga *et al*., 2012). In microbial ecology, however, it has been proposed that dispersal is rarely limiting as microbes can and do disperse widely, so environmental filtering is thought to be the main selective force determining differences in community membership, regardless of distance between sites (Astorga *et al*., 2012). Microbial communities, then, should only show strong distance decay relationships when sites that are closer together experience more similar environments (i.e., spatial autocorrelation in environmental conditions).

However, the relative strengths of dispersal limitation and environmental filtering in pond microbial communities may differ considerably from their hypothesized strengths in microbial communities in other ecosystems. First, dispersal limitation may be stronger than previously thought for pond-dwelling microbes. Indeed, high resolution sampling within ponds has revealed that the distance between sampling locations plays a critical role in shaping community composition, but that microbial dispersal can be limited at distances >20 m (Lear *et al*., 2014). On a larger scale, small lentic freshwater bodies like ponds are often more isolated than lotic freshwater environments, suggesting more limited dispersal among ponds compared to flowing aquatic systems (Shurin *et al*., 2009). This limited dispersal could generate steep distance decay relationships that may not be captured when sampling across broad geographic areas. Second, ponds and small lakes exhibit variable histories and contexts, from natural ecosystems to human-made water features stocked with non-native species, and placed in manicured parks, cemeteries, farms, and backyards. For these reasons, sites close by one another can be surrounded by vastly different landscapes (Hortal *et al*., 2014). Various factors within a pond can act as environmental filters shaping microbial communities, including turbidity, temperature, light pollution, nutrients, salinity, and several others (Premke *et al*., 2022; Huang *et al*., 2024; Karlicki *et al*., 2024; Niu *et al*., 2025). Thus, while we expect environmental filtering to play a substantial role in shaping biodiversity in small freshwater habitats, it is not clear how dispersal will interplay with environmental filters to shape distance decay relationships among ponds that may show weak spatial autocorrelation in environmental conditions.

Disentangling the main factors structuring microbial diversity, and whether they act either alone, in combination with, or in correlation with other landscape level factors remains a challenge in environmental microbiology (Huang *et al*., 2024). This goal is particularly challenging as some conditions are highly correlated with one another and the distribution of microbes across environmental and latitudinal gradients can follow linear or non-linear distribution patterns (Huang *et al*., 2024). To add complexity, drivers of community composition in freshwater systems may differ between the type of microbes (e.g., fungal vs bacterial), and microenvironments (e.g., pond muck vs pond water). Given the variety of factors that could promote variation in microbial membership across ponds, an outstanding question is whether ponds have a “core” microbiome or if these environments contain entirely unique taxa.

Here, we surveyed bacterial and fungal communities in water and muck from 39 freshwater ponds across the Eastern USA to characterize their biogeographical patterns. We tested for regional patterns of biodiversity, including a latitudinal diversity gradient and distance decay relationships. We also measured on-the-ground local environmental conditions in the pond and collected publicly available geospatial data on climate and land use and tested the relative importance of these factors in explaining variation in microbial community composition and richness. We used distance-based redundancy analysis followed by variance partitioning to tease apart various contributions of these variables in structuring microbial diversity. Understanding the major forces driving microbial diversity in freshwater systems will help us mitigate and manage these critical but often overlooked habitats in the face of environmental and anthropogenic change.

## Methods

### Water and muck sampling

We collected water and muck (organic detritus) during the spring and summer of 2022 in 8 states across the Eastern US (WI, NY, MI, ME, MS, GA, LA, NC). Ponds ranged in size from 6 to 25000 m^2^ (n = 39). While strict definitions vary, ponds are defined by being small (less than 2.5 hectares or 25000 m^2^), which generally allows for greater homogenous mixing through the water column. In comparison, small lakes (<4 hectares in area) stratify to form layers of temperature, oxygen, and nutrient levels, and undergo seasonal mixing regimes (Holgerson *et al*., 2022). For simplicity, we use the term pond to describe our sites, while acknowledging that others may define some as small lakes. At each site, 500 ml of water was collected from four locations around the outside of the pond and combined (2 L total). A filter line was primed by running water from the site through it without the sterivex attached. Pond water was passed through a series of cellulose filters to remove particulates [5 uM, followed by 3 or 1 uM] using a Masterflex peristaltic pump (Cole-Palmer North America, IL, USA). The water was then passed through a 0.22 uM sterivex filter, to collect free-living microbes, until the filter became saturated (the amount of filtered water was recorded = 578.2 ± 56.3 mL, mean ± SEM). The sterivex cartridge was wrapped in parafilm and stored on dry ice in the field before being transported to the University of Pittsburgh and stored at −80°C until DNA extraction. At each site, muck samples were also taken by placing a clean plastic straw into the surface layer of sediment (pond muck, ∼<3 inches deep) around the edge of the pond. The muck was stored in a clean container on dry ice before transport to the University of Pittsburgh and long-term storage at −80°C. Between sites, the filter line was rinsed in 70% ethanol, followed by deionized water and pre-filters were replaced.

At each of the four sampling locations around each pond, water quality was assessed using a YSI probe (YSI Incorporated, Ohio, USA) to measure dissolved oxygen, temperature, turbidity, chlorophyll, algae-phycocyanin (PC), salinity, and conductivity. Surface area of the water was estimated using a range finder to determine the relative width and length of the pond/lake (with surface area ∼ length x width). Muck pH was determined using a Hach H-Series H160 dual pH meter (Hach, CO, USA). Temperature and precipitation data from the day of sampling were obtained from the NASA Langley Research Center POWER Project funded through the NASA Earth Science Directorate Applied Science Program using the R package ‘nasapower’ (Sparks, 2018).

### Geospatial data

Climate and land-use data at the geographic coordinates for each pond were obtained from publicly available sources using R. Historical climate data for each pond were gathered from WorldClim 2.1 9 (Fick and Hijmans, 2017) using the ‘geodata’ package (Hijmans *et al*., 2023) to determine the average of monthly minimum temperatures (°C), monthly maximum temperatures (°C), and monthly precipitation (mm) from January 2012 to December 2021. Artificial night sky radiance (ALAN), or simulated zenith radiance in millicandelas per square meter (mcd/m^2^), was taken from the New World Atlas of Artificial Night Sky Brightness (Falchi *et al*., 2016). Land-use data were obtained from the USGS Annual National Land Cover Database (U.S. Geological Survey, 2024) using the ‘raster’ and ‘sp’ packages (Pebesma & Bivand, 2005; Hijmans *et al*., 2023), respectively). We calculated five land-use variables within a 500 m radius around each pond: the proportion of area classified as forest, wetlands, developed (to any extent), agricultural, and vegetation (grassland + shrubland), respectively.

### DNA extraction and sequencing

DNA was extracted from sterivex filters and muck samples using manufacturer instructions from a Powerfecal Pro Qiagen kit. For sterivex samples, the filters were removed from the casing manually using sterilized equipment and cut in half using ethanol and flame sterilized scissors. One half of the filter was used for DNA extraction with the other half reserved for microbial isolation and culturing (data not presented in this study). During extractions, muck and sterivex filters were added to homogenizing tubes prefilled with beads and homogenized for 10 minutes at room temperature before following directions for samples as described in the Powerfecal Pro instructions. The DNA concentration was quantified using nanodrop (Agilent, CA, USA) and then sent out for library preparation and sequencing on an Illumina Miseq platform at the University of Illinois, Chicago Genomics Research Core. The 16S rRNA gene was sequenced using 515F and 806R primers for the V4 region (read length 2 x 250). The ITS1 region of the ITS gene was amplified using ITS1F and ITS2R primers (read length 2 x 300). Our prefilter size likely filtered out larger fungal microbes from water samples, so we opted to only run fungal analysis on pond muck samples. The demultiplexed data was returned and processed using qiime2 and R (version 4.3.2) as described below.

### 16S rRNA data processing and analysis

Demultiplexed data was run through a qiime2 pipeline with additional analyses performed in R using phyloseq (v1.46.0; McMurdie & Holmes, 2013), microbiome (v1.24.0; Lahti & Shetty, 2018) decontam (v1.22.0; Davis *et al*., 2018), vegan (v2.6-8; Oksanen et al., 2013), microeco (Liu *et al*., 2021) and tidyverse (v2.0.0; Wickham *et al*., 2019) packages. Briefly, reads were assessed for quality before running them through Dada2 to trim, filter, denoise and pair ends (retained ∼76 ± 8% mean ± SD after dada2 – excluding samples with extremely low read counts). The reads were then classified using the greengenes2 classifier from the 2024.09 update (McDonald *et al*. 2023, Bokulich *et al*. 2018) and a rooted phylogenetic tree was constructed. The reads were filtered to include only ones identified as bacteria, and to remove chloroplast and mitochondrial DNA. Note that given the unexplored diversity of freshwater systems, we retained reads where taxonomic assignments of an amplicon sequence variant (ASV) was classified as “p___” indicating currently unnamed phyla. The resulting .qza data files were exported and the remaining analysis occurred in R. In R, reads were further processed to identify contamination based on negative kit controls using the prevalence method (threshold 0.5; decontam package). Several analyses were run on these data, as described in the statistical analysis section (taxa barplots, relative abundance analyses, and assessment of shared ASV across sites), using data filtered to exclude samples with low read counts (less than rarefaction level). Additionally, we assessed alpha diversity by calculating the number of observed ASVs in each sample, which is a proxy of richness (using data rarefied data without replacement to 3431 reads; 100 iterations). We also measured the Bray-Curtis dissimilarity, Jaccard and chi-squared distance metrics, which are all measures of beta diversity (using rarefied data; 100 iterations). Bray-Curtis dissimilarity is the main dissimilarity metric shown in the main text and is a measure of beta diversity that takes into account the abundance and presence/absence of ASVs. Next, we ran distanced based redundancy analysis (see statistical analysis) to identify environmental factors associated with variation in community composition. Finally, we assessed ASV rank abundance patterns across sample types on data rarefied without iteration since it was not built into the relevant vegan functions (radfit, specaccum).

### ITS1 analysis

Demultiplexed ITS1 data was also run through a qiime2 pipeline followed by analysis in R. The reads were assessed for quality and trimmed to avoid readthrough using cut-adapt. Next, the reads were run through Dada2 to trim, filter, denoise and pair ends into ASVs. Due to low quality in reverse reads, a large proportion of reads were removed at this step (retained ∼44 ± 9% mean ± SD after dada2 – not including samples with extremely low read counts). We also ran the analysis using only forward reads; however, this yielded poor taxonomic classification and greatly reduced estimates of biodiversity. Thus, we opted to move forward with paired ends. The reads were classified using a UNITE classifier for all eukaryotes with a dynamic classification standard, where species hypotheses are defined at the 97–99%, as recommended by fungal experts (Abarenkov et al., 2023) and the data was filtered to the phylum level as described above for 16S rRNA. The resulting data files were exported to R and processed for contamination. The data was then analyzed to generate taxa barplots, conduct relative abundance analyses (described below), and assess the amount of shared ASVs across sites, as described in the bacterial analysis. The data was also processed as described for the 16S rRNA to calculate alpha and beta diversity (using rarefied data to 2512 reads; 100 iterations). Finally, ASV abundance rank patterns were characterized, so that they could be compared to the patterns in pond bacterial samples (using rarefied data to 3431 reads to match bacterial data analyses).

Delineating fungal species based on similarity in ASVs is an ongoing research area, with some researchers clustering reads based on 97% operational taxonomic unit (OTU) thresholds as opposed to the full 100% ASV approach used here (Tedersoo *et al*., 2022). Additionally, we recognize that the presence of dead fungi and eDNA can lead to overestimates of fungal diversity that we cannot remove from analysis and ITS1 primers can introduce taxonomic bias due to an intron that is common in many fungi (Nilsson *et al*., 2019). We analyzed co-occurrence data based on (1) ASV and (2) taxonomic assignment (to the genus level; Figure S1), because analyzing at the level of ASV alone likely underestimates shared fungal membership across ponds and overestimates richness, as fungi often contain multiple ITS copies. However, limitations in reference databases make it so researchers currently recommend avoiding analyzing fungal taxonomy below the genus level (Nilsson *et al*., 2019). Decisions regarding data-processing in microbial ecology are known to produce divergent results, which makes it valuable to apply and compare different approaches to the data (Nearing *et al*., 2022).

### Statistical analysis

We performed multiple analyses across each of our sample types (muck, water) and microbial communities (bacteria, fungi) to (1) evaluate differences in community structure across sample types, (2) test for evidence of a core pond microbial community, (3) determine biogeographical patterns, and (4) identify environmental factors that explain variation in microbial community richness and composition. To start, we evaluated the evenness of the community by plotting rank abundance curves, where ASVs were ordered based on abundance of reads across samples and plotted against relative ASV abundance. Here, steeper curves indicate less even communities. The data were standardized across sample types by rarefying to 3431 reads per sample and randomly sampling to the same number of total samples (n = 30) across types (muck fungal, water bacterial, muck bacterial). Curves were fitted using radfit in the vegan package, where the best fit model (between zipf, zipf-mandelbrot, lognormal, null, and preemption) was selected and plotted (Shoemaker *et al*., 2017). Second, we compared the global community richness across all sites, sample types and sample sizes by plotting the log10 of the number of unique ASVs against the number of ponds included in the estimate of richness. Here, estimates were determined using speccacum function with the random method and 100 permutations. Third, we looked for shared taxa across the majority of sites, indicating a core microbial community (Neu *et al*., 2021) by plotting the frequency of occurrence for each ASV against the log10 of the relative abundance of the ASV using code adapted from Burns et al., [2016]. We also generated taxa barplots which can be found in the supplement (Figure S2, S3).

Next, we examined whether there was evidence of a latitudinal diversity gradient by comparing a null model to a linear model relating alpha diversity to latitude with or without pond area as a covariate using Bayesian information criterion (BIC). The best model was the one with the lowest BIC, although all models with ΔBIC < 7 of the best model were considered (see associated R markdown for details). Then, we tested for evidence of a distance decay relationship between microbial community similarity (equal to 1 - Bray-Curtis dissimilarity) and distance between sites. We fit commonly reported linear and non-linear distance decay models to the data and compared model fits by BIC among these models and a null model.

To examine environmental drivers of beta diversity, we started by assessing correlations between environmental parameters. Variables that were highly correlated (>0.7 correlation coefficient) included (1) water temperature in the pond with precipitation the day of sampling and climate variables of temperature, (2) phycocyanin and chlorophyll, (3) development and light at night (ALAN). We opted to include only one of each of these variables in further analysis (pond water temperature, chlorophyll, development), while noting that the high correlation between these variables and the others precludes us from determining which is the primary driver of variation in community composition. We then generated a global distance-based redundancy analysis (dbRDA) model of Bray-Curtis dissimilarity that included environmental variables (e.g., oxygen, temperature, pH, land use variables) not highly correlated with one another. All explanatory variables were scaled to ensure they were equally weighted in the analysis (although we tested with and without doing scaling and it did not affect the conclusions). To determine the best model, selection was performed using ordistep() function in R.

Following dbRDA analysis, we ran variance partitioning analysis (varpart function, vegan package) to determine how much variation in the dissimilarity matrix (for Bray-Curtis, Jaccard, and chi-squared) was associated with the best-fit model for environmental variables versus the distance between sites (using geodesic distances to account the curvature of the earth in the distance estimation; geodist package, v0.1.0; Padgham, Mark; Sumner, 2025).

Connections between environmental factors (scaled as in dbRDA analysis) and alpha diversity were also assessed using a global linear model fit to the data. Model fits were compared by BIC using the dredge function in R. There were several models with ΔBIC < 7 of the best-fit model. We moved forward with the top model in all cases, as many variables were only present in a few of the alternative models, while explanatory variables in the best-fit model were included in several of the models with ΔBIC < 7, and thus, contained the strongest evidence for inclusion. This result means that other factors may explain variation in alpha diversity across ponds, but that the evidence in this dataset was weaker than those present in the best-fit model.

To explore the relationship between environmental variables and specific taxa, we performed ANCOM-BC2 (using AMCOMBC package; Lin & Peddada, 2020, 2024) and the microeco package). We evaluated relative abundance at the family level against environmental variables that were found to be significant in either the beta diversity or alpha diversity analysis. For water bacterial samples, this included temperature, oxygen, and log10(conductivity). For muck bacterial samples this included temperature and log10(conductivity). For muck fungal samples this included temperature and log10(conductivity). Briefly, the unrarefied data was filtered to remove samples with low read counts (less than 3431 for bacteria and 2512 for fungi). Only families present in >30% of samples with an average of >0.1% abundance across samples were included in the analysis.

## Results

### Abundance distributions

ASVs for water bacterial and muck fungal communities followed common rank abundance patterns, where some ASV had high abundance, while most were rare (Figure 1a). However, muck bacterial communities had a broader and shallower rank abundance curve compared to the other sample types, indicating greater evenness (Figure 1a). The top model depended on the microbial community, with water bacterial communities being best represented by a Zipf model, muck bacterial communities by a Zipf-Mandelbrot, and fungal communities by a lognormal model (see associated R Markdown file).

**Figure 1.**
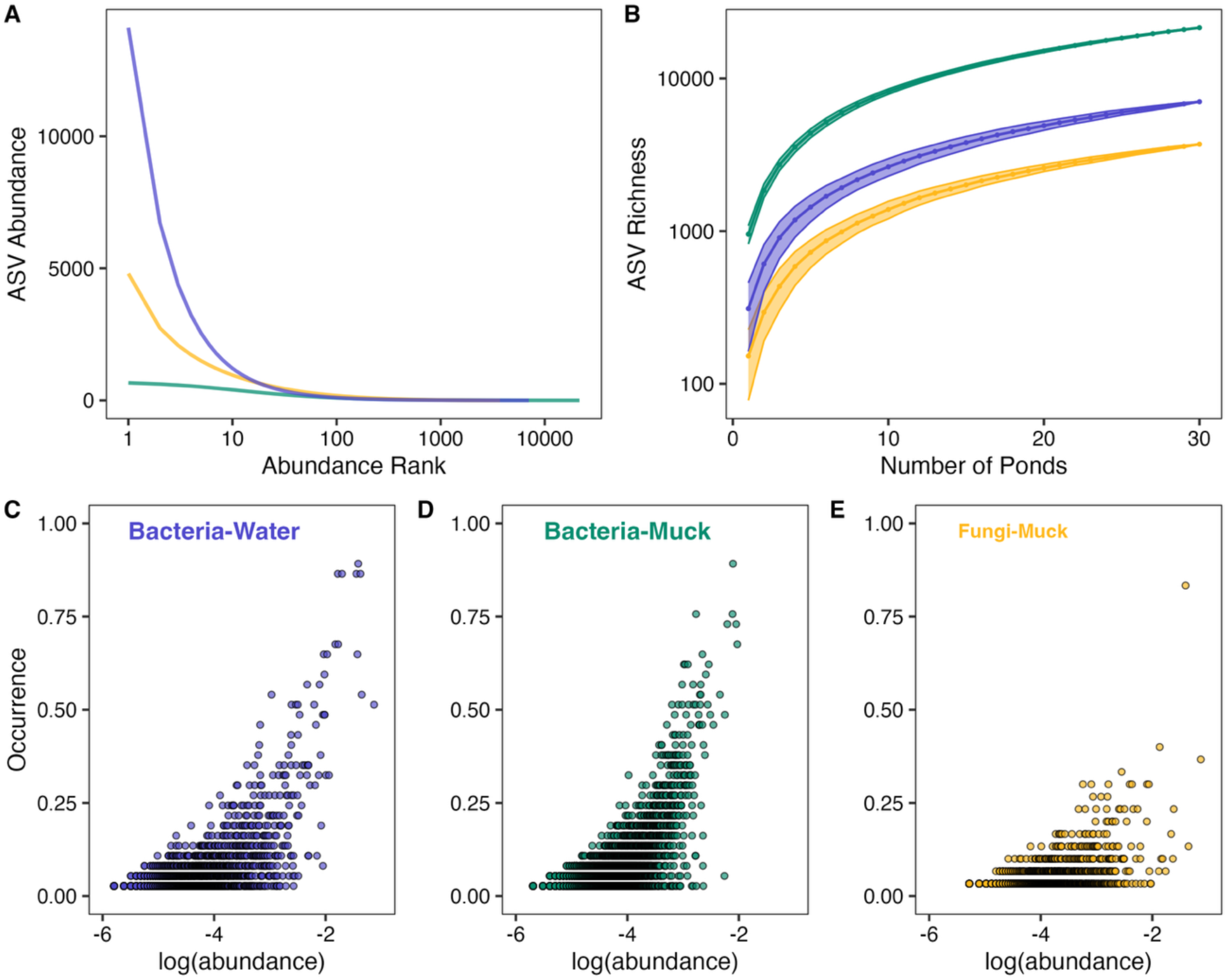
Diversity dynamics of pond microbial communities. Plots show the ASV abundance distributions for bacterial communities in muck (green) and water (blue), and fungal communities in muck (yellow). Plot (A) shows rank abundance curves where ASVs were ranked based on abundance across all reads and plotted against ASV rank abundance. For (B) The log10 of the number of unique ASV (richness) is plotted against the number of ponds included in the estimate of richness. In (C–E), each point represents a different ASV, where the occurrence (proportion of ponds containing that ASV) is plotted against the log10 of the relative abundance of the ASV. Note that plots (A) and (B) are on rarefied data, while (C–E) is on unrarefied data, to improve detection of shared ASV across ponds.

The majority of ASV were found in <25% of ponds, with fungal communities containing the least number of shared ASVs (almost all ASVs were found at <50% of ponds), but also the lowest richness (Figure 1C–E). However, fungal occurrence across ponds increased at the genus level (Figure S1). In water bacterial communities, five ASV were found across >80% of sites and the most abundant phyla were Pseudomonadota, Actinomycetota, and Bacteroidota. These ASVs were in the genus *Planktophila* (phylum = Actinomycetota), *Sediminibacterium* (phylum = Bacteroidota), *Aquirufa_904070* (phylum = Bacteroidota), and *Polynucleobacter* (phylum = Pseudomonadota). A total of 2122 bacterial genera from 1060 families were identified in pond water across all sites. Pond muck was similarly dominated by Pseudomonadota, Actinomycetota, and Bacteroidota, though the communities differed in the relative abundance of classes within these phyla (Figure S2). In pond muck, only one ASV was found in >80% of ponds, which was in the family *Xanthobacteraceae* (phylum = Pseudomonadota). A total of 2800 bacterial genera from 1291 families were found in pond muck across all sites. Muck communities also contained high amounts of the phyla Acidobacteriota, Planctomycetota, and Chloroflexota, which were scarce in water. In muck fungal communities, Ascomycota made up >50% of reads on average. Basidiomycota was the second most abundant division followed by Chytridiomycota. These divisions were also highly abundant across pond sites, representing >75% of relative abundance on average (Figure S3 shows taxa bar plots at the class level).

### Latitudinal diversity gradient

There was moderate evidence of a latitudinal diversity gradient in muck fungal communities, with fewer observed ASVs from ponds at higher latitudes (Figure 2, ΔBIC between null and top model = 4.68; p-value = 0.006, adjusted R2 = 0.199). For bacterial communities in muck and water, there was limited evidence of a latitudinal gradient, as the best-fit models were the null. Pond size was not a significant covariate across any of our sample types.

**Figure 2.**
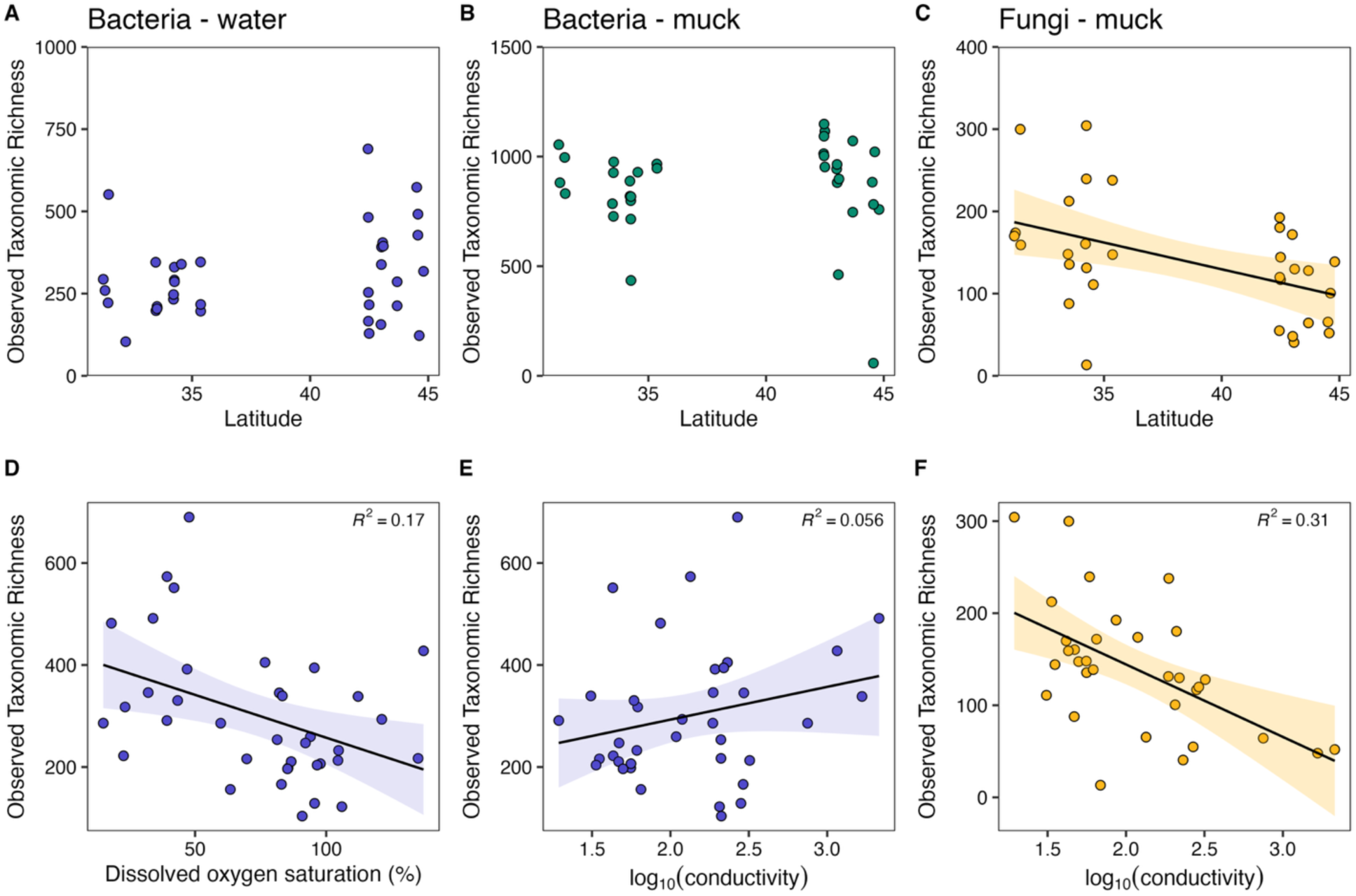
Assessing evidence of latitudinal diversity gradients in pond microbial communities. (A) bacterial communities in pond water, (B) bacterial communities in pond muck, (C) fungal communities in pond muck. Where applicable, the best fit model is overlaid on the plot. (D-F) shows relationships between environmental variables and alpha diversity metrics. Plots showing significant environmental drivers related to observed taxonomic richness (number of ASVs) in (D and E) water bacteria and (F) muck fungal communities in ponds.

### Environmental drivers of richness

Microbial richness was associated with environmental filters for water bacterial and muck fungal communities, although exact relationships differed across sample types (Figure 2). For water bacterial communities, the best-fit model included dissolved oxygen and log(conductivity) (adjusted R^2^ = 0.238). For muck fungal communities, the best-fit model incorporated log(conductivity) (R^2^ = 0.31, adjusted R^2^ = 0.286). Several other models were within 7 ΔBIC of the best-fit model, although they frequently contained the environmental variables represented in the best-fit model, with more scarce representation of additional factors, indicating weak evidence that these factors contributed substantially to variation in ASV richness across sites. The best model was the null for muck bacterial communities.

### Distance decay relationships

In pond muck and water, the best-fit model for both fungal and bacterial communities was a simple linear model between community similarity and distance between sites (Figure 3). Slope values ranged from −2.5e^−5^ in the bacteria in water, to −2.6e^−5^ in the bacteria in muck, and −1.5e^−5^ in the fungi in muck. Here, a steeper slope indicates a more rapid decrease in similarity across space, indicating greater species turnover.

**Figure 3.**
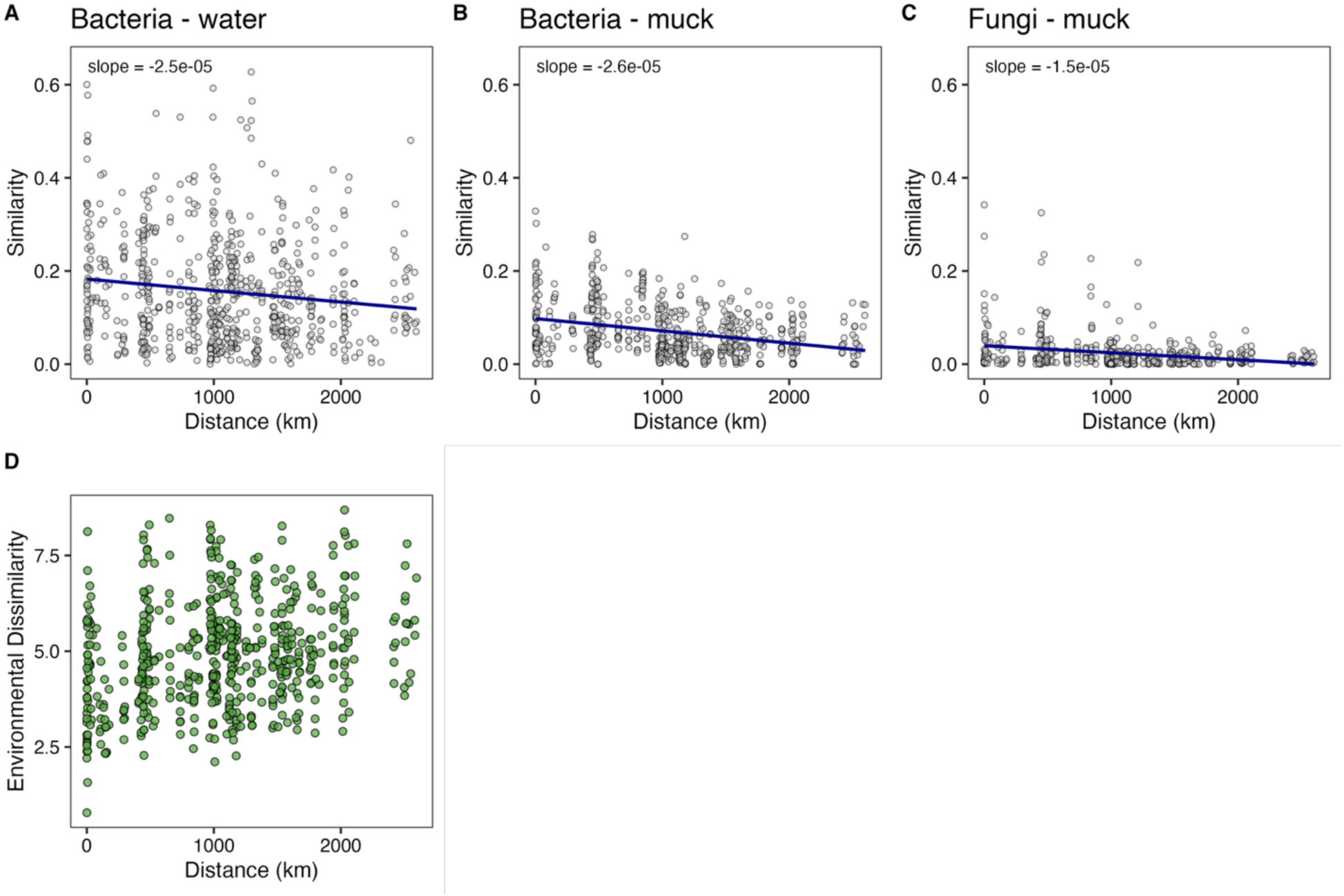
Assessing evidence of distance decay relationships in pond microbial community composition. (A) bacterial communities in pond water, (B) bacterial communities in pond muck, and (C) fungal communities in pond muck. The best fit model is overlaid, which was a simple linear model for distance between sites and Bray-Curtis similarity in microbial communities. (D) is the relationship between the environmental dissimilarity between two ponds and the distance between those ponds. Each point on the graph represents a comparison between two ponds. The y-axis is the geodesic distance between those two sites (accounts for Earth’s curvature in distance estimate), while the x-axis shows the Euclidean distance between the environmental parameters at the two sites. A mantel test (Pearson method) revealed a weak but significant relationship between the two (Mantel statistic R = 0.246).

### Environmental drivers of community structure

Environmental and landscape filters explained portions of microbial community compositions, though the main explanatory factors varied across sample types (Figure 4). For bacteria in pond water, the best-fit dbRDA model included information about water temperature, surrounding forest cover, dissolved oxygen in the water, chlorophyll content, and surface area of the pond (Figure 4). The model explained ∼27% of the variation in the data (adjusted R^2^ =0.147), with dbRDA 1 and dbRDA 2 capturing ∼11% and ∼5%, respectively. For muck bacterial communities, the best-fit model included log(conductivity), water temperature, precipitation, and surrounding forest cover, accounting for ∼18% of variation (adjusted R^2^ = 0.070), with dbRDA 1 covering ∼7% of total variation. For muck fungal communities, the best-fit model explained ∼9% of variation in the data (adjusted R^2^ = 0.0248) and included precipitation and water temperature, with dbRDA 1 accounting for ∼5% of the total variation.

**Figure 4.**
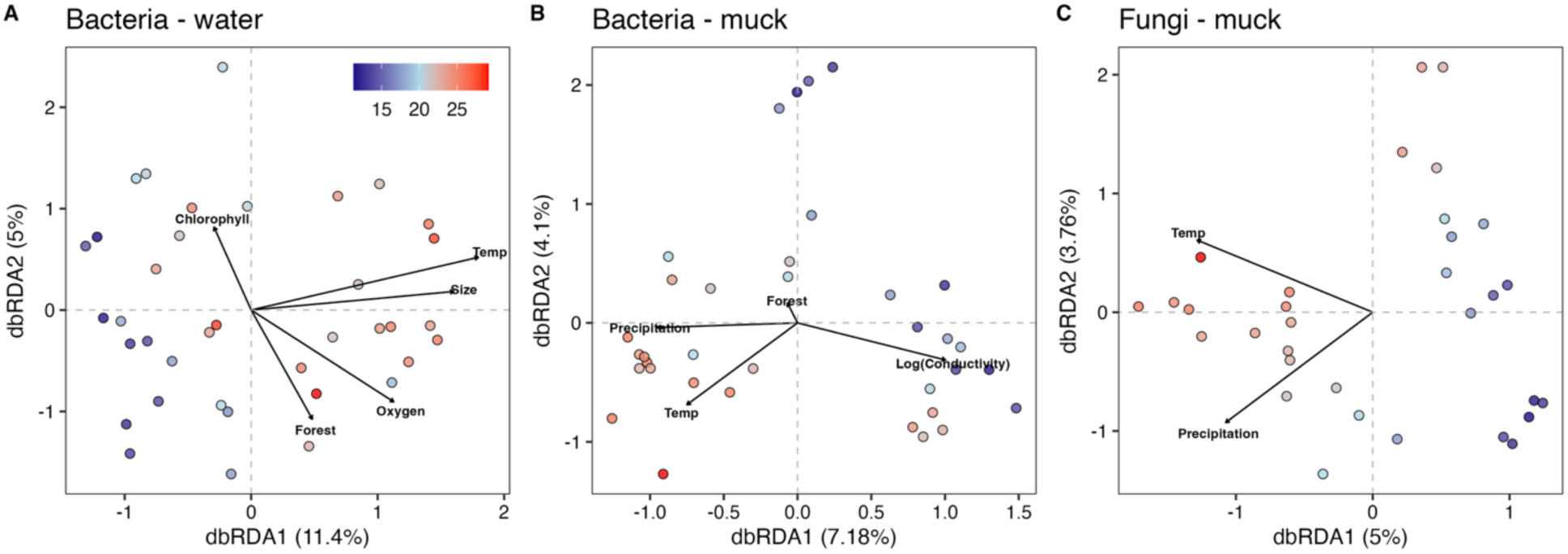
Relationships between environmental variables and beta diversity metrics. Results of distance-based redundancy analysis (dbRDA) using Bray-Curtis dissimilarity. Panels show the best model for dbRDA of pond microbial communities as they relate to environmental and landscape parameters. The color of the points indicates the pond water temperature (°C) during sampling, which was found to be a significant factor explaining variation in beta diversity in bacterial and fungal communities.

We used variance partitioning to compare the effects of distance and the environmental and land use filters found to be significant in the dbRDA analysis (water temperature, surrounding forest cover, dissolved oxygen in the water, chlorophyll content, and surface area). Results revealed that local factors had stronger impacts in structuring pond water communities than distance between sites (Table 1). In pond muck, very little variation in community composition could be uniquely explained by environmental factors or distance between sites (Table 1).

**Table 1.**
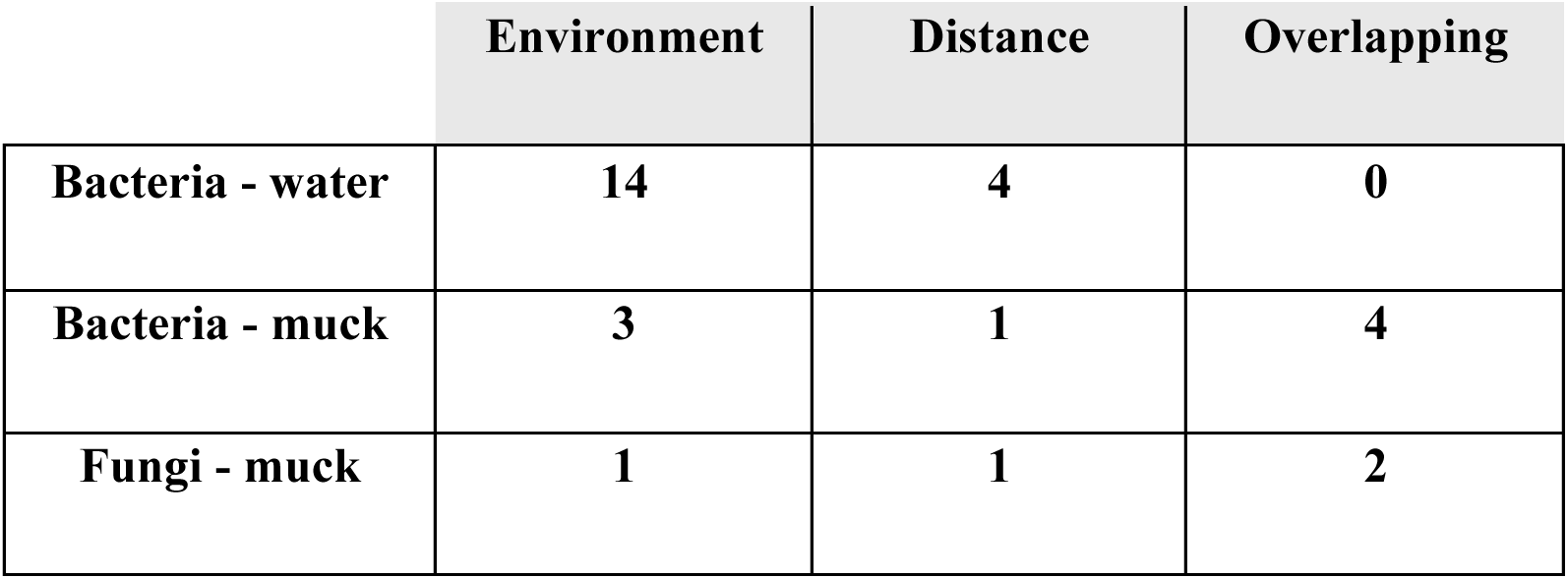
Percent of variation explained by distance and environmental factors. We used variance partitioning to compare the effects of distance and the environmental and land use filters (water temperature, surrounding forest cover, dissolved oxygen in the water, chlorophyll content, and surface area).

|  | Environment | Distance | Overlapping |
| --- | --- | --- | --- |
| **Bacteria - water** | **14** | **4** | **0** |
| **Bacteria - muck** | **3** | **1** | **4** |
| **Fungi - muck** | **1** | **1** | **2** |

ANCOM-BC2 revealed a number of bacterial families that were significantly associated with environmental drivers (Table S1). In muck bacterial communities, Xanthobacteraceae [phylum Pseudomonadota, also known as Proteobacteria] and uncultured family SbA1 [phylum Acidobacteriota] were associated with conductivity (log fold-change = −1.73 and −2.53, respectively). Water bacterial communities also contained one family, Nostocaceae [phylum Cyanobacteriota] whose relative abundance was related to conductivity (log fold-change = - 4.78). Additionally, the relative abundance of three families, Methylomonadaceae [phylum Pseudomonadota], Nostocaceae [phylum Cyanobacteriota], Casimicrobiaceae [Pseudomonadota], were related to dissolved oxygen saturation in the water (Table S1; log fold-change = −0.03, 0.07, 0.07, respectively). There were no significant relationships between environmental drivers and specific families in the muck fungal communities.

## Discussion

Here we sampled pond water and muck from across the Eastern USA to explore factors shaping microbial biodiversity, as significant discoveries in ecology and evolution have stemmed from observations on geographic variation in biodiversity (Gaston K.J., 2000). In particular, there is rising interest in evaluating whether patterns commonly observed in animals and plants also apply to microbial communities (Xu *et al*., 2020; Härer & Rennison, 2023). Small and highly variable environments, like freshwater ponds and tiny lakes, are unique, island-like ecosystems that harbor rich biodiversity of plants and animals (Davies *et al*., 2008), but are also commonly found in human-altered spaces, including cities, parks, farms, and backyards (Hill *et al*., 2017). Due to their distinctive traits, ponds are valuable systems for evaluating whether microbial communities follow known biogeographical patterns in animals, investigating factors that shape community assembly, and evaluating how human activity influences these dynamics.

### I. What microbes live in ponds?

Pond bacterial communities were dominated by phyla, such as or Pseudomonadota (also known as Proteobacteria), Actinomycetota, and Bacteroidota, which are often found in freshwater environments, both lentic and lotic (Jezbera *et al*., 2009; Shannon *et al*., 2023; Huang *et al*., 2024; Karlicki *et al*., 2024; Rohwer *et al*., 2024; Wang *et al*., 2024; Niu *et al*., 2025). For example, the most abundant phyla in muck and water, Pseudomonadota, was similarly abundant in sediment from freshwater streams, lakes, ponds, and rivers in a country-wide sampling effort in Germany (Premke *et al*., 2022), wastewater sludge (Wu *et al*., 2019). Pseudomonadota were also highly abundant in sampling of global freshwater and marine shipping ports (Ghannam *et al*., 2020) and from inland water samples collected across China (Wang *et al*., 2024). When comparing bacterial abundance patterns across all sites, muck bacteria exhibited a shallower species rank abundance curve, suggesting higher evenness, while water bacteria had a much steeper curve, with a few ASVs dominating the global community.

In muck and water, bacteria displayed similar abundance-occurrence relationships, where there were a small number of commonly occurring ASVs. Unsurprisingly, the handful of ASVs found in the greatest number of ponds (>80%), belonged to widely observed bacterial classes, including Actinomycetes, Bacteroidia, Gammaproteobacteria, and Alphaproteobacteria. Across the three different microbial communities surveyed, muck fungal communities contained the lowest amount of shared ASVs across ponds, with >99% of ASVs only being found in <25% of sites. However, at higher taxonomic levels, muck fungal communities were composed of widely distributed fungal divisions that are known to live in sediments and soils, such as Ascomycota, Basidiomycota, and Chytridiomycota (Tedersoo *et al*., 2014; Tian *et al*., 2018). Though pond microbial communities were similar at higher levels (division/phyla, class), the low number of shared ASV across sites points to high rates of turnover driven by deterministic or stochastic processes, such as dispersal limitations, environmental filtering, and diversification (Xu *et al*., 2020; Dickey *et al*., 2021). This could result in communities maintaining similar ecological services across ponds, despite high variability in community membership at the ASV level due to functional redundancy (Ramond et al., 2025).

### II. What factors shape richness of pond microbial communities?

Next, we investigated factors affecting local microbial diversity. Latitudinal diversity gradients are one of the most widely observed patterns in biogeography (Hillebrand, 2004). Elucidating the mechanisms driving latitudinal diversity gradients is an active area of research, with some proposing that gradients are caused by abiotic and biotic variables, like temperature, which covary with latitude (Hillebrand, 2004; Huang *et al*., 2024). We found evidence of a latitudinal diversity gradient in muck fungal communities, but not in bacterial communities from water or muck. These findings contrast with other large scale sampling efforts using standardized methodology that have found latitudinal gradients in bacteria in inland waters (Wang *et al*., 2024) and paddy field soils (Huang *et al*., 2024), and some soil fungal communities (Tedersoo *et al*., 2014). However, the strength of these relationships and generality of latitudinal diversity gradients are inconsistent across studies, with meta analyses finding no evidence of latitudinal diversity gradients in freshwater systems, or among bacteria or fungi (Kinlock *et al*., 2018).

When investigating specific environmental variables, microbial richness was negatively related to dissolved oxygen saturation and positively related to conductivity in water bacterial communities. These results add to growing evidence that oxygen is tightly linked to microbial richness in aquatic systems (Beman & Carolan, 2013; Spietz *et al*., 2015), though others have observed unimodal rather than linear relationships between oxygen and richness, where diversity declines at exceptionally low oxygen (Beman & Carolan, 2013). As oxygen levels can fluctuate both daily and seasonally, it will be interesting for future work to examine how richness varies temporally within the same pond. Interestingly, the richness of muck bacterial communities was not related to any environmental or land-use variables, demonstrating that the abiotic factors in the water do not appear to influence the overall alpha diversity of the muck. In contrast, the richness of muck fungal communities was negatively correlated with conductivity, but surprisingly, not to temperature or muck pH, which are known drivers of fungal diversity in terrestrial soil (Islam *et al*., 2020; Mikryukov *et al*., 2023) and freshwater lake sediment (Tian *et al*., 2018). Muck fungal communities in ponds may have distinct sensitivity to abiotic conditions, though it is also important to consider methodological differences (e.g., region of ITS gene used, sampling protocol, statistical analyses) that could influence results across studies (Tedersoo *et al*., 2022).

Ultimately, sensitivity to environmental drivers could affect resilience within the community to future environmental perturbation and their vulnerability to human-alterations of the system. Oxygen, for example, differs based on surrounding floral diversity and management practices, such as the presence of pond aeration systems. Conductivity is also influenced by human activities, where water salinity is higher in areas that use road salt or coastal regions vulnerable to seawater intrusion (Cañedo-Argüelles *et al*., 2019; Reid *et al*., 2019). Though surrounding agriculture was not significantly related to microbial richness here, run-off from agricultural sites may still lead to eutrophication or contaminants in ponds which could influence biodiversity on scales not captured here or could lead to harmful algal blooms which are known to impact water oxygen levels (Reid *et al*., 2019). During decision making, it is important to consider links between management and restoration on microbial diversity, through changes in the abiotic environment.

### III. What drives variation in pond microbial community composition across broad geographic areas?

Here, we observed that pond microorganisms exhibit relatively low community similarity with modest distance decay relationships. Many macro- and microorganisms exhibit strong exponential decays, though substantial variation in the decay slope has also been observed across taxa and studies (Graco-Roza *et al*., 2022). Our findings align with those from aquaculture ponds, where bacterial communities have also exhibited weak distance decay relationships and steeper slopes in sediment communities compared to water (Niu et al., 2025). Additionally, soil from lakeshore wetlands showed low overall similarity across fungal and bacterial communities, but with slightly steeper slopes than observed here (Xu *et al*., 2024).

Distance decay relationships in macro-organisms are often attributed to dispersal limitations and filtering via environmental gradients (Nekola & White, 1999; Astorga *et al*., 2012). While it has been proposed that microbial biodiversity follows a rule of “*microbes are everywhere and the environment selects*” (Soininen, 2012), microbes also seem to be subject to dispersal limitations in ponds (Lear *et al*., 2014). We found that pond environmental conditions had low spatial autocorrelation with distance, suggesting that these habitats exhibit a high amount of environmental variability even within small geographic areas. Given the island like features of ponds as isolated, lentic water bodies, dispersal may be more limited in these environments compared to lotic and larger bodies of water, especially across large geographic areas. Muck may be especially dispersal-limited due to its higher water content and distinct qualities compared to terrestrial soil, which could constrain the ability for microbes to survive outside of the pond and disperse to other sites, even at short distances, though this requires further investigation.

Several abiotic factors, such as temperature and dissolved oxygen, were significantly related to microbial community composition across ponds, though the overall amount of variation explained by these filters was relatively low. Additionally, oxygen and conductivity were associated with the relative abundance of several bacterial families. Water temperature was a universal driver of community composition across sample types, aligning with prior work across a variety of habitats, including terrestrial soils, wastewaters, marshes, and inland waters (Martiny *et al*., 2011; Wu *et al*., 2019; Mikryukov *et al*., 2023; Wang *et al*., 2024). We also evaluated whether land-use could explain community differences, as this is often used as a proxy for other water quality parameters, like nutrient pollution and toxicant exposure (Cheng *et al*., 2022). Interestingly, forest cover was associated with variation in bacterial community structure, while development, agriculture, and vegetation (grassland + shrubland) were not. Forest cover could affect composition through multiple mechanisms, including by altering the light and temperature environment in the ponds. In Jyrkänkallio-Mikkola et al., [2017], bacterial community composition in Finland streams was also related to surrounding forest composition, but not agriculture. The unexplained variation may be attributable to environmental filters known to influence microbes that were not directly measured here (e.g., nutrients, pollutants; Niu et al., 2025) or differences in stochastic processes (e.g., drift, priority effects; Langenheder & Lindström, 2019).

There were a few families of bacteria that were significantly associated with conductivity or oxygen in the water. One of these, Nostocaceae, is a family of photosynthetic cyanobacteria that was related to conductivity and oxygen. Relative abundance of this family was positively related to oxygen in the water, potentially because they are oxygen-producing. These types of cyanobacteria can form colonies and bacterial mats that act as harmful algal blooms (Hentschke *et al*., 2025). In our dataset this family was present in low average abundance and sparsely distributed across samples (in only ∼30% of the water samples). Additionally, the abundance of the aerobic methanotroph family, *Methylomonadaceae,* was negatively related to oxygen, but not conductivity, despite showing a strong negative relationship with salinity during a sampling of sediment from freshwater, brackish, and saline lakes in Tibet (Deng *et al*., 2024). Though they are considered aerobic, it is also thought that this family may be resilient to low oxygen conditions, which may explain the relatively weak relationship between oxygen and their relative abundance observed here (Deng *et al*., 2024). Of note, two families of bacteria were also strongly negatively related to muck conductivity, including an uncultured family of Acidobacteriota, SbA1, that are commonly found in soil (Mcreynolds *et al*., 2025).

## Future Work

While we did not find a substantial impact of development or agricultural land use here, these factors may play a role depending on more nuanced variation in land management, including the specific type of development, treatments performed on the water and landscape topography and drainage, which would not be captured in medium resolution Earth-observing satellite datasets. For example, land managers often aerate ponds or add commercial products to the water, like algicides, cleaners, and probiotics. These manipulations are generally unregulated and/or undocumented. Thus, the scale and breadth with which we can measure anthropogenic effects may limit our ability to determine their role. Stochastic factors, like drift, unpredictable disturbances and priority effects (Debray *et al*., 2022), may also shape lentic freshwater biodiversity, though this requires further investigation. Many ponds are ephemeral or semi-permanent, meaning that they may dry out during part of the year and then partially or fully reform during the wet season, leading to more stochastic variation. Additionally, seasonal variation in environmental conditions can contribute to temporal changes in microbial biodiversity that will be important to consider when evaluating drivers across spatial scales (Rohwer *et al*., 2024). We encourage future work studying the impacts of various human manipulations on microbial biodiversity across space and time.

Future work may also address whether differences in community composition translate to functional characteristics of the community. Variation in microbial community composition does not necessarily generate functional differences, especially given patterns of functional redundancy across environments (Lear *et al*., 2014; Grossart *et al*., 2020; Graco-Roza *et al*., 2022). Approaches such as metagenomics, meta-transcriptomics, and metabolomic analyses will help reveal how environmental filters may be altering microbial physiology or selecting for different strains (Grossart *et al*., 2020). Indeed, rapid and repeated evolution within a bacterial species can occur in freshwater environments (Rohwer *et al*., 2024). For example, long-term metagenomic inventories of a lake in Madison, WI revealed that bacterial strains within the same species fluctuated seasonally and repeatedly year after year, in cases where the relative abundance of those species remained the same (Rohwer *et al*., 2024). These nuanced changes are not resolvable through 16S rRNA sequencing but point to avenues for future work.

## Conclusion

Our findings reveal distinct biogeographical patterns of pond microorganisms. Despite coming from the same source environments, biogeographical trends were conditional on the type of microbe and microenvironment. Specifically, we found evidence of a latitudinal diversity gradient only in pond muck fungal communities. We found microbial richness was positively related to conductivity in bacteria from water but negatively related to richness in muck fungi. We also observed several consistent patterns across all three microbial communities. All sample types showed weaker distance decay relationships compared to lotic freshwater systems and macro-organisms, suggesting shared ecological factors governing turnover and dispersal across ponds. Temperature was also a universal driver of variation in community composition across microenvironments, though its importance differed depending on the microbe and microenvironment. While environmental conditions explained more variation in microbial community composition than distance between sites, both factors collectively still explained minimal overall variation, suggesting a larger role for stochastic processes or environmental variables not evaluated here. Collectively, the limited spatial autocorrelation in environmental conditions, coupled with the lack of connectivity among ponds offers a useful framework for investigating localized drivers of microbial diversity. Future work should explore how this underlying variation impacts the function of these communities, such as contributions to water quality, decomposition, nutrient cycling, and host-microbe associations within these critically important habitats.

## Data and Code Availability

The raw sequence data and relevant metadata have been deposited in the NCBI Sequence Read Archive (SRA) under BioProject accession number PRJNA1378205.

## Acknowledgements

We are especially grateful to Cory Duckworth for assistance in developing methods and collecting samples.

## AI disclosure

We used AI-based tools (ChatGPT, grammarly) to generate some basic suggestions for code structure and copy-editing improvements. However, all outputs were reviewed, edited, and verified by the authors.

