## Supplemental figures for "Exploring local and regional drivers of microbial biodiversity across freshwater ponds"

**Table S1. Significant results from Ancom-BC2 analysis**

| Sample Type | Kingdom | Phylum | Class | Order | Family | Environmental variable | Log fold-change | Adjusted p-value |
|---|---|---|---|---|---|---|---|---|
| Muck bacterial | Bacteria | Pseudomonadota | Alphaproteobacteria | Rhizobiales_505101 | Xanthobacteraceae | log10(conductivity) | -1.73 | 0.0416 |
| Muck bacterial | Bacteria | Acidobacteriota | Terriglobia | Terriglobales | SbA1 | log10(conductivity) | -2.53 | 0.008 |
| Water bacterial | Bacteria | Pseudomonadota | Gammaproteobacteria | Methylococcales | Methylomonadaceae | Dissolved oxygen saturation (%) | -0.03 | 0.0088 |
| Water bacterial | Bacteria | Cyanobacteriota | NA | NA | NA | log10(conductivity) | -3.86 | 0.0212 |
| Water bacterial | Bacteria | Cyanobacteriota | Cyanobacteriia | Cyanobacteriales | Nostocaceae | Dissolved oxygen saturation (%) | 0.07 | 0 |
| Water bacterial | Bacteria | Cyanobacteriota | Cyanobacteriia | Cyanobacteriales | Nostocaceae | log10(conductivity) | -4.78 | 0.0219 |
| Water bacterial | Bacteria | Pseudomonadota | Gammaproteobacteria | Burkholderiales | Casimicrobiaceae | Dissolved oxygen saturation (%) | 0.07 | 8.00E-04 |

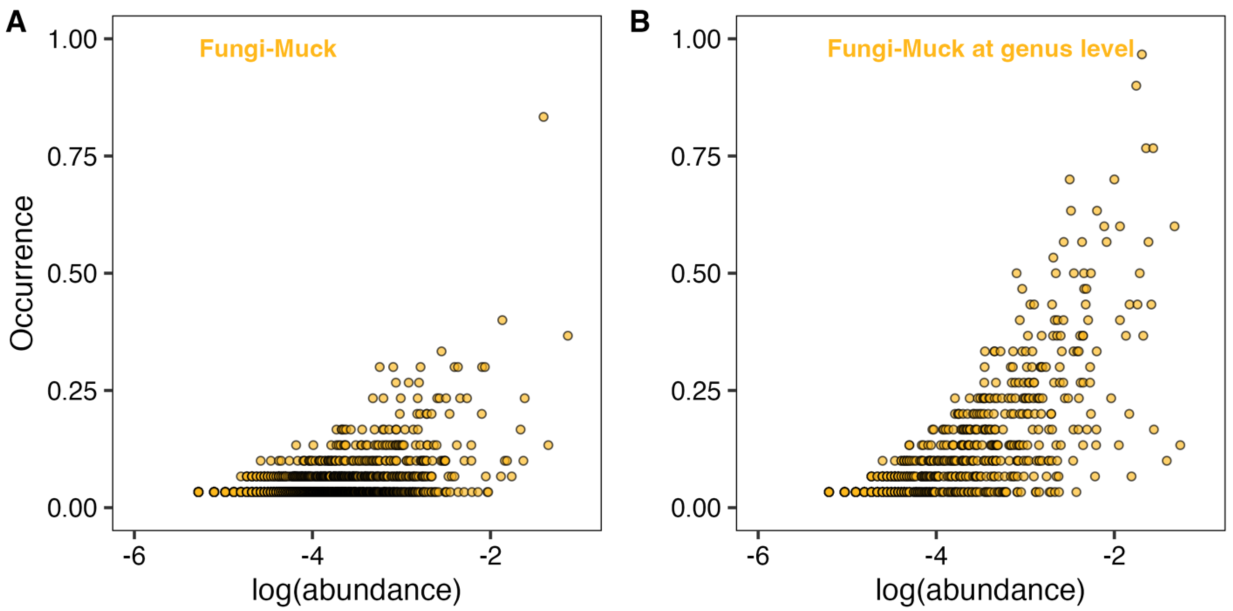


**Figure S1.** Comparison of abundance-occurrence patterns for pond muck fungal communities at the (A) ASV vs (B) genus level. Each point represents a different (A) ASV or (B) genus, where the occurrence (percent of ponds containing that taxa) is plotted against the log10 of the relative abundance of the taxa.

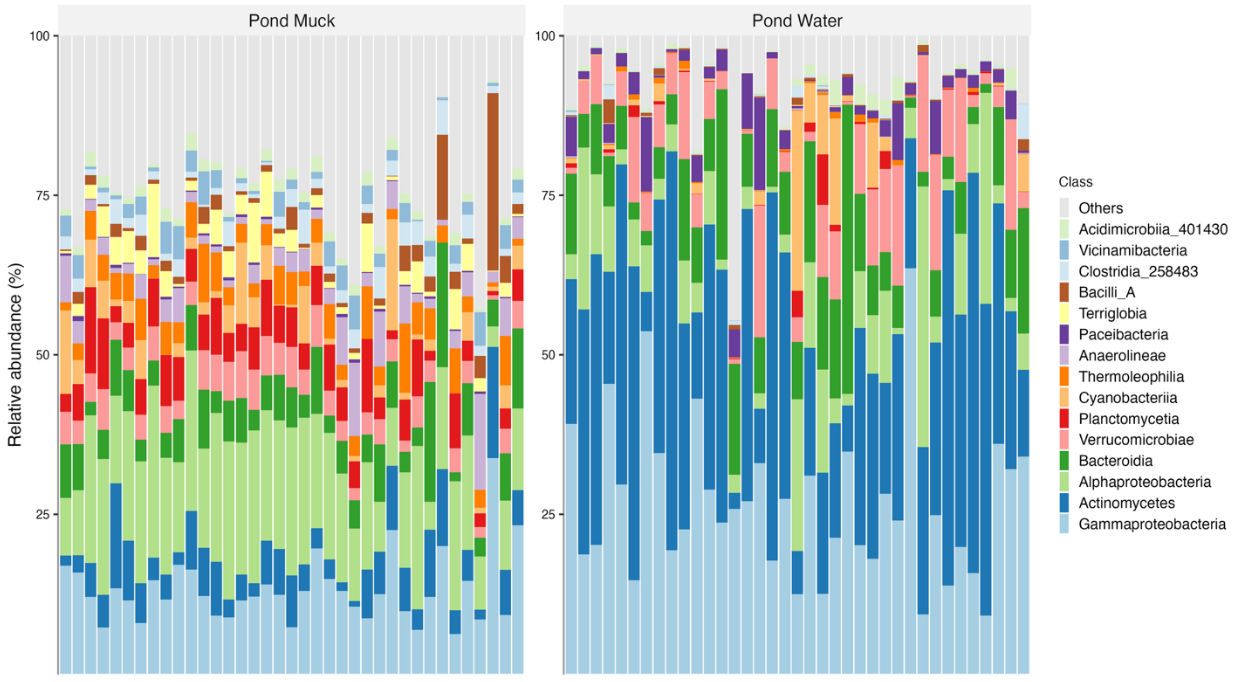


**Figure S2.** Taxa bar plots showing relative abundance of the top 15 most abundant classes found in pond bacterial communities. Each bar represents the community found in a different pond, with muck samples on the left panel and water samples on the right panel.

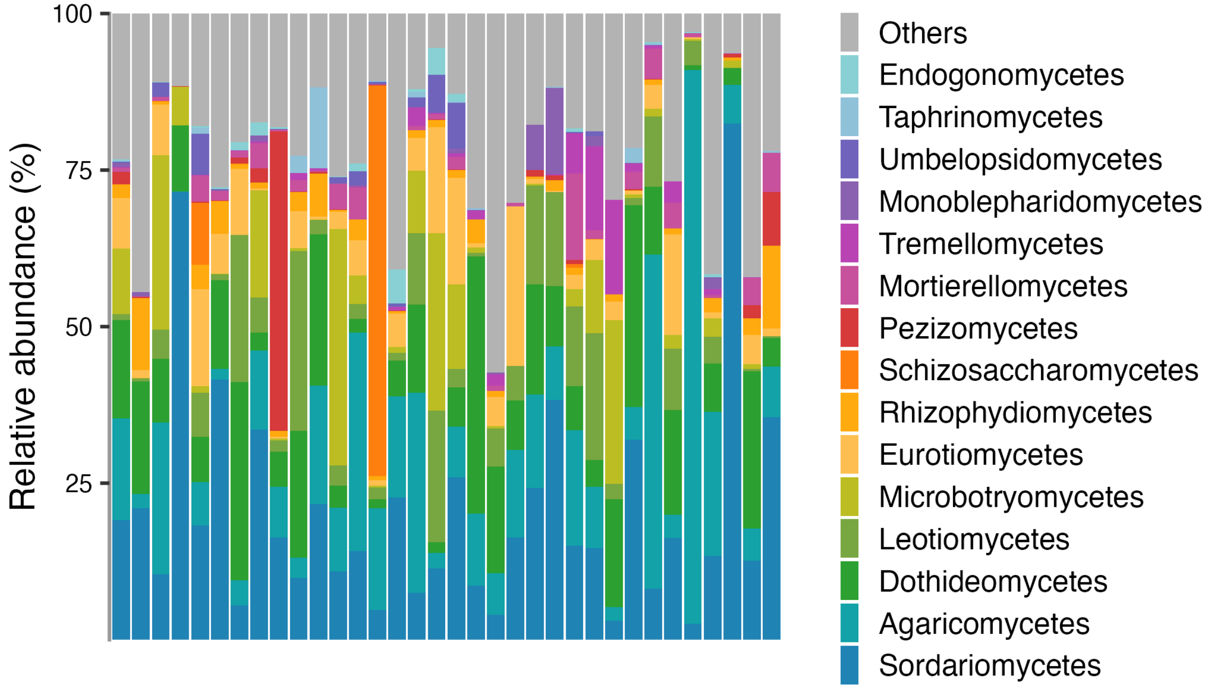


**Figure S3.** Taxa bar plots showing relative abundance of the top 15 most abundant classes found in pond fungal communities. Each bar represents the community found in a different pond.

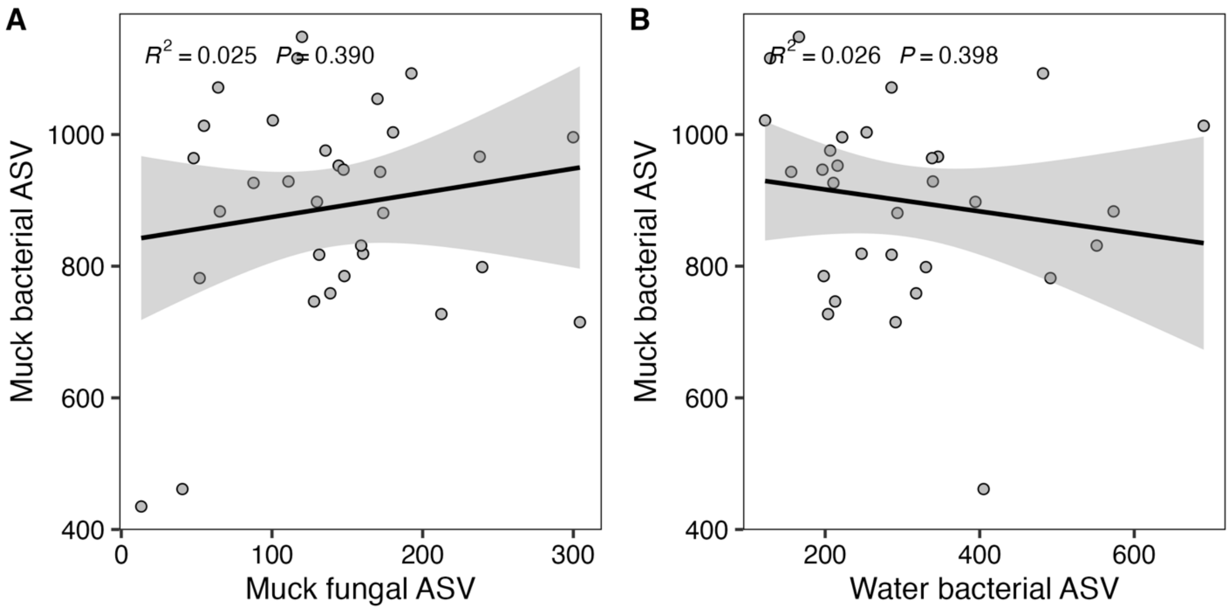


**Figure S4.** Comparison of ASV richness across different sample types within the same pond. (A) shows the relationship between muck and water bacterial richness and (B) shows muck bacterial and muck fungal communities. Lines show linear fits with confidence intervals from geom_line() in ggplot, with $R^2$ values reported on the plot.
